# 2P-FLIM unveils time-dependent metabolic shifts during osteogenic differentiation with a key role of lactate to fuel osteogenesis via glutaminolysis identified

**DOI:** 10.1101/2023.01.13.523892

**Authors:** Nuno GB Neto, Meenakshi Suku, David A. Hoey, Michael G. Monaghan

**Affiliations:** Department of Mechanical, Manufacturing and Biomedical Engineering, Trinity College Dublin, Dublin, Ireland; CURAM SFI Research Centre for Medical Devices, National University of Ireland, Galway Ireland; Advanced materials for Bioengineering Research (AMBER), centre, Trinity College Dublin and Royal College of Surgeons in Ireland, Dublin, Ireland

## Abstract

Human mesenchymal stem cells (hMSCs) fuel discrete biosynthetic pathways to multiply and differentiate into specific cell lineages; with undifferentiated hMSCs showing reliance on glycolysis. hMSCs differentiating towards an osteogenic phenotype rely on oxidative phosphorylation as an energy source. Here, the metabolic profile of hMSCs was profiled during osteogenic differentiation over 14 days using a non-invasive live-cell imaging platform- two-photon fluorescence lifetime imaging microscopy (2P-FLIM) which images and measures NADH fluorescence. During osteogenesis, we observe a higher dependence on oxidative phosphorylation for cellular energy; concomitant with an increased reliance on anabolic pathways. We validated this metabolic profile using qPCR and extracellular metabolite analysis and observed a higher reliance on glutaminolysis in the earlier time-points of osteogenic differentiation. Based on the results obtained, we sought to promote glutaminolysis further during osteogenic differentiation. An indirect method of promoting glutaminolysis was explored so as to not impact cellular differentiation. As Lactate has been shown to promote glutamine uptake via c-Myc activation triggering expression of glutamine transmembrane transporters and glutaminase 1; we chose to increase extracellular lactate concentrations to drive increased glutaminolysis rates leading to higher levels of mineral deposition and osteogenic gene expression. Lactate supplementation of osteogenic medium also promoted upregulation of lactate metabolism and increased the expression of transmembrane cellular lactate transporters. Higher rates of lactate dehydrogenase gene expression coupled with higher NADH fluorescence intensity demonstrate a conversion of lactate to pyruvate. During this conversion, NADH is formed by the reverse enzymatic reaction of lactate dehydrogenase resulting in increased NADH fluorescence intensity. In order to evaluate the importance of glutaminolysis and lactate metabolism in osteogenic differentiation, these metabolic pathways were shut down using BPTES and α-CHC respectively which led to reduced hMSC mineralisation. In summary, we demonstrate that hMSCs osteogenic differentiation has a temporal metabolic profile and shift that is observed as early as day 3 of cell culture. Osteogenic differentiation was demonstrated to be directly dependent on OxPhos and on glutaminolysis and validated using biochemical assays. Furthermore, extracellular lactate is an essential metabolite to ensure osteogenic differentiation as a metabolic fuel and signalling molecule to promote glutaminolysis. These findings have significant impact in generating potent approaches towards bone tissue engineering *in vitro* and *in vivo* by engaging directly with metabolite driven osteogenesis.

## Introduction

hMSCs differentiation towards an osteogenic, adipogenic or chondrogenic lineage is accompanied by changes in phenotype, gene expression, protein secretion and cellular metabolism that occur in a time-dependent manner [1]. Metabolic regulations of hMSCs differentiation are still not fully understood [2] but in general it is known that stromal cells rely on glycolysis for energy when undifferentiated. Such a profile that is dependent on glycolysis facilitates a quick turnover of energy whilst producing biomolecules required for cellular proliferation [3]. hMSCs continue to utilize glycolysis as the main source of energy during chondrogenic differentiation [4]. However, there is a metabolic switch to oxidative phosphorylation (OxPhos) during hMSCs differentiation towards adipogenic or osteogenic lineages [5, 6]. In addition, it has been shown that an increase in oxygen consumption and increase of mitochondrial activity are important for osteogenic differentiation of hMSCs [5, 7]. Most recently, glutamine metabolism has been shown to have an important role during hMSCs differentiation into osteogenic lineages [8].

Two-photon fluorescence lifetime imaging (2P-FLIM) is a powerful technique for non-invasive probing and monitoring of cellular metabolism. This technique is based on the fluorescence properties of nicotinamide adenine dinucleotide (NAD(P)H) and flavin adenine dinucleotide (FAD). NAD(P)H fluorescence lifetime consists of short- and long-lifetime components corresponding, respectively, to free and protein-bound forms [9]. NAD(P)H is an important metabolic co-factor that drives ATP production and other anaplerotic metabolic pathways. NAD(P)H drives glycolysis in the cytoplasm and oxidative phosphorylation (OxPhos) in the mitochondria. Therefore, longer fluorescence lifetimes are associated with higher ratios of protein-bound NAD(P)H revealing a higher dependence of OxPhos as a source of ATP. NAD(P)H fluorescence signal, especially protein-bound NAD(P)H (τ1), can be used to infer quantitatively in NADPH concentrations [10]. NADPH has an important role in lipid, amino acid and nucleotide biosynthetic pathways as well reactive oxygen species (ROS) protection [11]. FAD^+^ is also an important metabolic co-factor for OxPhos. Specifically, FAD^+^ is a proton acceptor during the Krebs cycle and a proton donor on the electron transport chain (ETC) [12]. The ratios of the fluorescence intensities of FAD and NAD(P)H (FAD/NAD(P)H) are referred to as optical redox ratio and it can be used to estimate the metabolic profile of cells or tissues [13]. 2P-FLIM has been used as a technique to investigate cellular metabolism in a variety of cells, tissues and organoids [14–16]. After imaging, the sample can be still used for further metabolic, biochemical or histological validation [17]. In addition, it is possible to include machine learning models to the data analysis to further understand the impact of metabolic treatments [18].

hMSCs metabolic probing during differentiation using 2P-FLIM has been previously reported yet several of these studies are limited to direct observation of metabolic changes at a specific time-point without other biochemical validation such as evaluation of extracellular fluxes, measuring metabolites concentration or observing mitochondrial shape or biogenesis. In addition, considerations regarding hMSCs source and media formulation are also not made [19–21]. One study reported by Meleshina et al., applied 2P-FLIM to monitor hMSCs metabolic profile while in osteogenic differentiation culture for 21 days [19]. An increase of OxPhos was verified in hMSCs as early as day 7 of cell culture. Furthermore, the authors suggest that reliance on OxPhos osteogenic differentiation of hMSCs increases when cellular proliferation is less pronounced. However, this work does not detail the complete media formulation used to induce osteogenic differentiation in the study. Therefore, it is difficult to determine if cells were differentiated in supraphysiological conditions impacting both hMSCs differentiation and cellular metabolism. Moreover, this work did not probe the cellular metabolic profile of differentiated hMSCs using standard biochemical techniques. Media formulation in particular, are one important consideration overlooked when investigating hMSCs bioenergetics, possibly influencing cellular metabolism due to supraphysiological levels of nutrients [5, 7, 22].

In this study, we profile hMSCs osteogenic differentiation using 2P-FLIM to uncover metabolic shifts. Afterwards, the metabolic profiles observed during hMSCs osteogenic differentiation were validated using extracellular metabolite quantitation and gene expression analysis (Figure 1). Another aim of this study was to improve osteogenic differentiation by promoting specific anaplerotic pathways. A study by Yu et al., demonstrated that glutaminolysis is crucial in ensuring adequate osteogenic differentiation of hMSCs [8]. Due to the importance of glutamine and glutaminolysis to osteogenic differentiation, we decided to drive further osteogenic differentiation by upregulating glutaminolysis. We hypothesize that supplementing osteogenic differentiation medium with exogenous lactate promotes higher glutaminolysis rates and higher yields of cellular differentiation. Perez-Escuredo et al., have demonstrated that lactate uptake by monocarboxylate transporter 1 (MCT1) promotes glutamine uptake and metabolism in oxidative cancer cells by c-Myc activation [23]. In that manner, an increased uptake of glutamine and glutaminolysis rate culminating in alpha-ketoglutarate production will possibly increase osteogenic cell differentiation as demonstrated by increasing mineral deposition. Our results show that 2P-FLIM derived τ_avg_ values determined an OxPhos profile during osteogenic differentiation. However, increased OxPhos is associated with higher ORR parameters due to a higher mitochondrial electron transport chain (ETC) activity. This increase in ETC activity, converts NADH to NAD^+^ reducing NADH fluorescence intensity and increases FAD^+^ concentration resulting in increased FAD^+^ fluorescence intensity. Instead, 2P-FLIM ORR results show a decrease in ORR during osteogenic differentiation. This occurs due to higher NADH concentrations resulting in an increase of NADH fluorescence intensity. Therefore, during osteogenic differentiation, hMSCs rely on OxPhos and on anaplerotic pathways that promote the generation of NADH.

**Figure 1.**
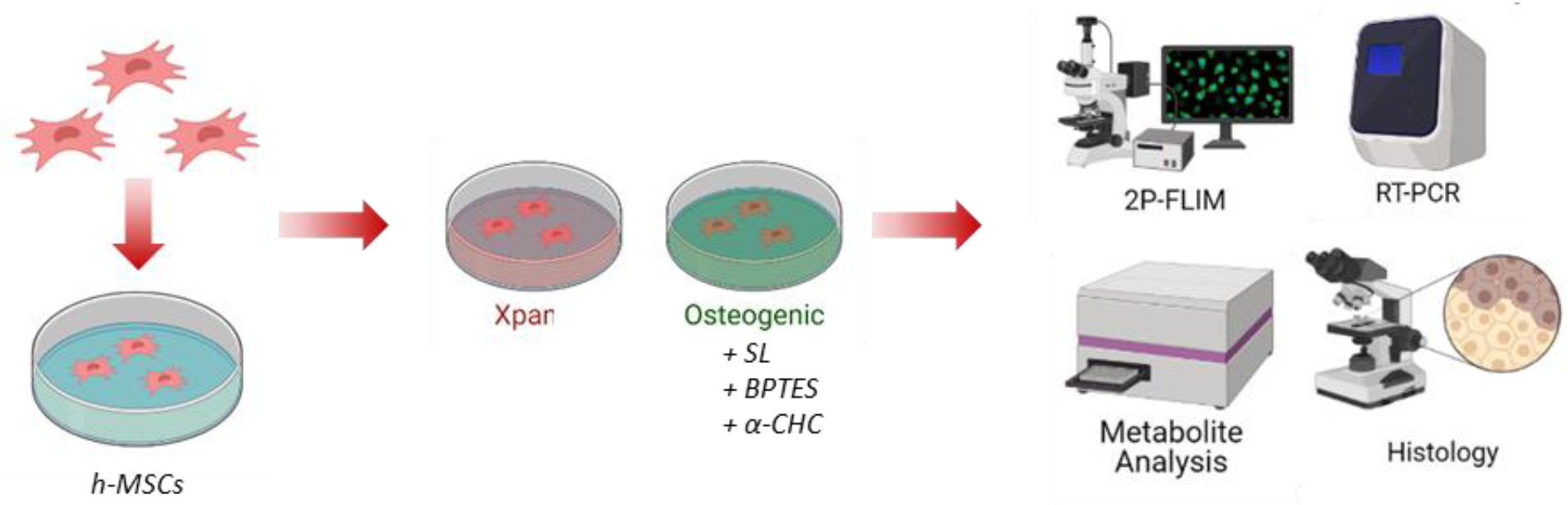
Experimental Design. SL-sodium lactate; BPTES - Bis-2-(5-phenylacetamido-1,3,4-thiadiazol-2-yl)ethyl sulphide; α-CHC - α-Cyano-4-hydroxycinnamic. Image created using Biorender^®^

## Results

### Osteogenic differentiation involves Oxidative Phosphorylation in a time-dependent manner

hMSCs were cultured in Xpan and Osteo + media for 14 days. On day 7 and day 14 of culture, hMSCs were fixed using paraformaldehyde (PFA) and stained with alizarin red to verify mineral deposition and osteogenic differentiation (Figure 2A). As expected, after 14 days of incubation a noticeably higher mineral deposition in Osteo + medium conditions were observed, associated with increased osteogenic differentiation (figure 2A, Supplemental figure 1). Osteogenic gene expression of hMSCs was quantified with qPCR, after 14 days of culture in either Xpan or Osteo + media (Figure 2 B, C, D). A statistically significant increase in the gene expression of (2.3-fold increase) alkaline phosphatase (ALPL) and (2.19-fold increase) prostaglandin-endoperoxide synthase 2 (PTGS2) was observed in hMSCs when cultured in Osteo + medium (Figure 2B). The expressions of the remaining osteogenic genes were found to be not statistically significant between Osteo + and Xpan media conditions. PCA visualization of osteogenic gene expression revealed a statistically significant segregation between hMSCs incubated in Xpan or Osteo + cell culture medium with p-value<0.05 and F-value higher than critical F-value (Figure 2C, Supplemental Table 2). A z-score heatmap based on osteogenic gene expression shows similar results with a higher expression of ALPL and PTGS2 in Osteo + conditions after 14 days of incubation (Figure 2C).

**Figure 2.**
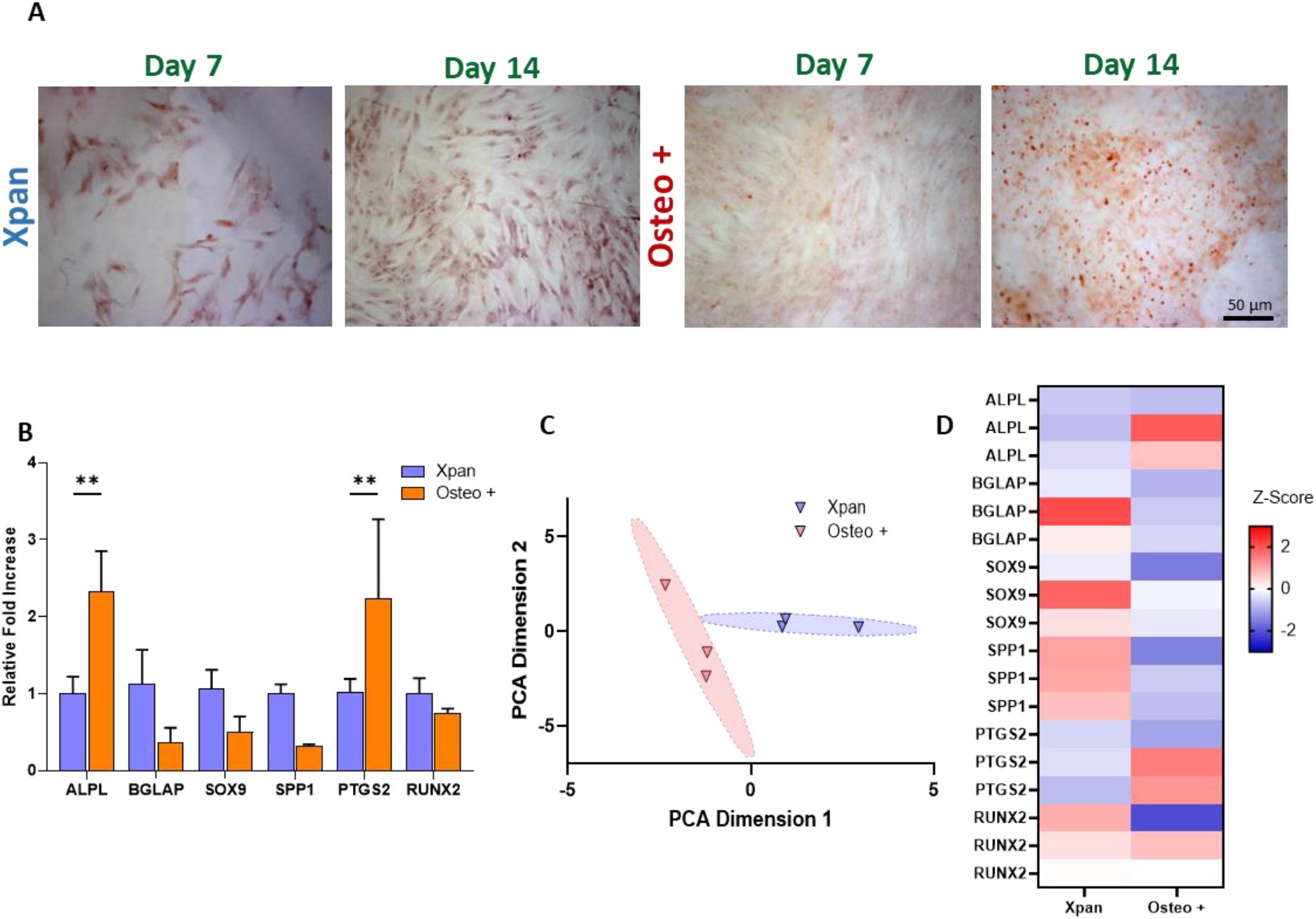
Validation of osteogenic differentiation of hMSCs incubated in Osteo + or Xpan cell culture medium for 14 days. A) Alizarin red staining of hMSCs after 14 days in contact with Osteo + or Xpan medium. B) hMSCs gene expression of osteogenic markers after 14 days of cell culture. C) Z-score heatmap of hMSCs gene expression of osteogenic gene markers after 14 days of cell culture. D) PCA distribution and of hMSCs osteogenic gene expression after 14 days of cell culture. *p-value≤0.05, **p-value≤0.01, ***p-values≤0.001, ****p-value≤0.0001 N≥3

2P-FLIM of NAD(P)H and FAD^+^ revealed a time-dependent metabolic shift with hMSCs in osteogenic medium (Figure 3). On day 0, no significant differences in τ_avg_, τ_1_ or ORR values were observed between hMSCs cultured in Xpan or Osteo + media (Figure 3 B, C, D). On day 3 of cell culture, a significantly higher (1.097 ± 0.04 ns versus 1.167 ± 0.004 ns) τ_avg_ and (2.683 ± 0.027 ns versus 2.800 ± 0.039 ns) τ_1_ values concomitant with lower (0.724 ± 0.020 versus 0.615 ± 0.021) ORR values (Figure 3 B, C, D) could be seen. The trends observed on day 3 were found to become more accentuated with longer term cultures. NAD(P)H fluorescence τ_avg_ (1.167 ± 0.004 ns at day 3, versus 1.236 ± 0.010 ns at day 7 and 1.316 ± 0.011 ns at day 14) and τ_1_ (2.800 ± 0.039 ns at day 3, versus 2.979 ± 0.053 ns at day 7 and 3.146 ± 0.015 ns at day 14) was found to be higher in Osteo + treated cells while in Xpan conditions, these fluorescence variables had constant values. Similarly, ORR values continued to decrease in Osteo + conditions after day 3 (0.615 ± 0.021 ns at day 3, versus 0.534 ± 0.029 ns at day 7 and 0.585 ± 0.012 ns at day 14) while hMSCs in Xpan medium had stable ORR values (Figure 3 B, C, D).

**Figure 3.**
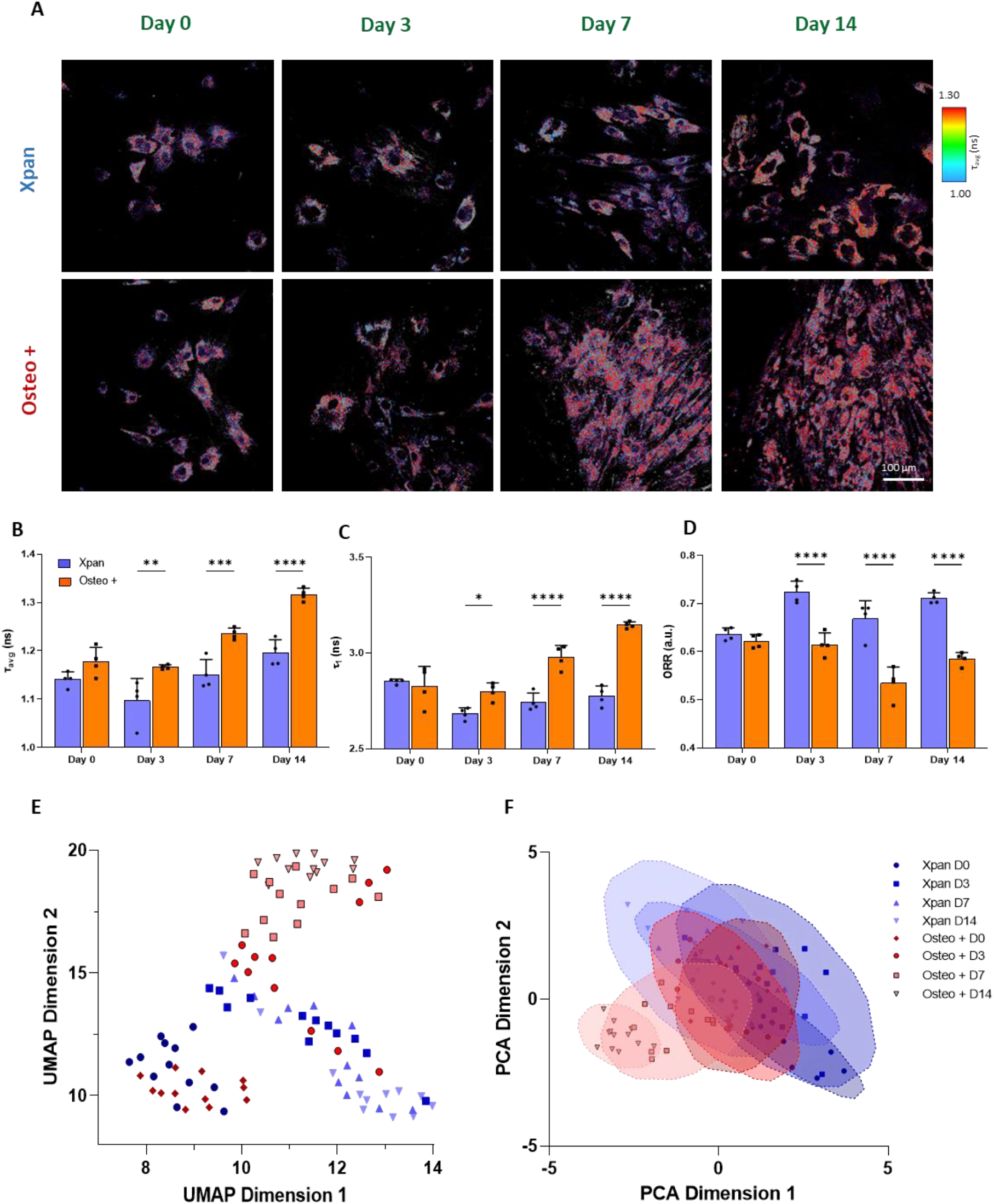
Metabolic profiling of hMSCs incubated in Osteo + or Xpan cell culture medium for 14 days. **A) 2P-FLIM imaging of hMSCs at day 0, 3, 7 and 14 of cell culture incubation.** B) NAD(P)H τ_avg_ measurement of hMSCs at day 0, 3, 7 and 14 of cell culture. C) NAD(P)H protein-bound lifetime τ_1_ at day 0, 3, 7 and 14 of cell culture. D) Optical Redox Ratio (ORR) of hMSCs at day 0, 3, 7 and 14 of incubation in cell culture medium. E) UMAP distribution of 2P-FLIM NAD(P)H and FAD^+^ fluorescence lifetimes and intensity. F) PCA distribution of 2P-FLIM NAD(P)H and FAD^+^ fluorescence lifetimes and intensity. *p-value≤0.05, **p-value≤0.01, ***p-values≤0.001, ****p-value≤0.0001 N≥3

UMAP and PCA were used as exploratory data visualisation tools to infer the metabolic profile of hMSCs undergoing osteogenic differentiation at several time-points during cell culture (Figure 3 E, F). All 2P-FLIM NAD(P)H and ORR variables were used to generate the UMAP and PCA plots. UMAP revealed that for later time-points, such as day 7 and day 14, there is a strong separation between hMSCs cultured in either Osteo + or Xpan media (Figure 3 E). For earlier time-points, there was no segregation between hMSCs cultured in both cell culture medium conditions (Figure 3F, Supplemental table 2). When applying PCA, there is a clear and statistically significant segregation of cells treated with Osteo + media at day 7 and day 14 compared with hMSCs cultured in Xpan media. (Figure 3 F, Supplemental table 2).

To further validate the time-dependent metabolic shifts during osteogenic differentiation, cell culture media was collected during media exchanges at days 2, 5, 8 and 11. Glucose, lactate and glutamine concentrations were measured using colorimetric assays (Figure 4 A, B, C). On assessing the glucose consumption, a statistically significant increase was observed (2.42-fold increase) when cells were cultured in Xpan medium at all time-points (Figure 2A). Additionally an increasing trend in lactate secretion was noted in both Xpan and Osteo + treated hMSCs during the incubation period. On day 11, there was a statistically significant higher lactate secretion in hMSCs cultured in Xpan medium (4.660 ± 0.384 mM) compared with hMSCs in Osteo + (3.295 ± 0.406 mM) culture conditions (Figure 2B). Lastly, a significantly higher glutamine consumption was found on day 2 of hMSCs in Osteo + (1.417 ± 0.017 mM) compared to Xpan (1.000 ± 0.041 mM) cell culture conditions. For the remaining time-points there was a decreasing trend of glutamine consumption for both cell culture media conditions (0.01-fold for Osteo + and 0.32-fold for Xpan) (Figure 4C).

**Figure 4.**
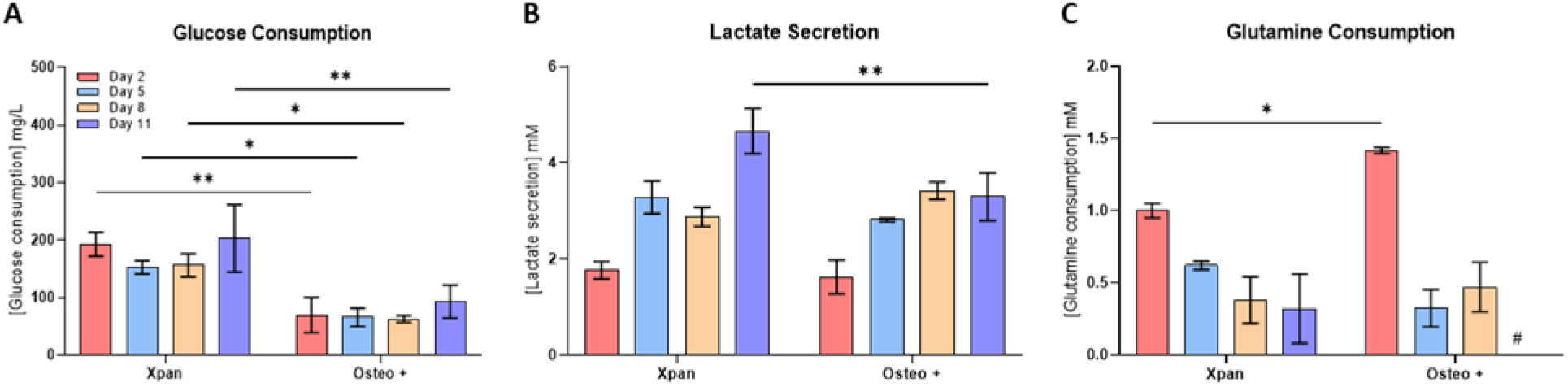
Extracellular metabolite consumption rates of hMSCs cultured in Osteo + or Xpan cell culture medium for 14 days. A) hMSCs glucose consumption at day 2, 5, 8 and 11 of cell culture. B) Lactate secretion at day 2, 5, 8 and 11 of cell culture. C) hMSCs glutamine consumption at day 2, 5, 8 and 11 of cell culture. #no glutamine consumption, *p-value≤0.05, **p-value≤0.01, ***p-values≤0.001, ****p-value≤0.0001 N≥3.

### Lactate supplementation promotes higher mineral deposition and impacts metabolism

hMSCs osteogenic differentiation promoted a metabolic shift towards OxPhos and increased dependence on anabolic pathways such as glutaminolysis. Here, increased dependence of glutaminolysis and anabolic pathways was induced by adding exogenous lactate to osteogenic medium. Lactate supplementation of osteogenic medium yielded a higher mineral deposition after 14 days of culture with a higher amount of alizarin red staining when compared with Osteo + medium (Figure 5A). Osteogenic genes expression was measured by qPCR in hMSCs incubated in Osteo +, Xpan and Lactate supplemented osteogenic medium (Figure 5 B, C, D). We observed a statistically significant increase in SRY-Box Transcription Factor 9 (SOX9) (2.7-fold increase) and Osteopontin (SPP1) gene expression (15.47-fold increase) in hMSCs when osteogenic medium was supplemented with lactate compared to Osteo + and Xpan cell culture medium (Figure 5 B, D). RUNX2 gene expression is also slightly higher in lactate supplemented osteogenic medium compared with Xpan (1.39-fold increase) and Osteo + (1.87-fold increase) incubation medium (Figure 5 B, C). PCA distribution plot of cell culture conditions of hMSCs at day 14 shows a statistically significant segregation between hMSCs incubated in Osteo + Lact, Osteo + and Xpan based on the expression of their osteogenic genes with p-value<0.05 and calculated f-value higher than critical f-value (Figure 5C, Supplemental table 2). ALPL, BGLAP, PTGS2 have similar gene expression values in hMSCs when incubated in osteogenic medium with and without lactate supplementation, as observed in the bar plot and the z-score heatmap (Figure 5 B, D).

**Figure 5.**
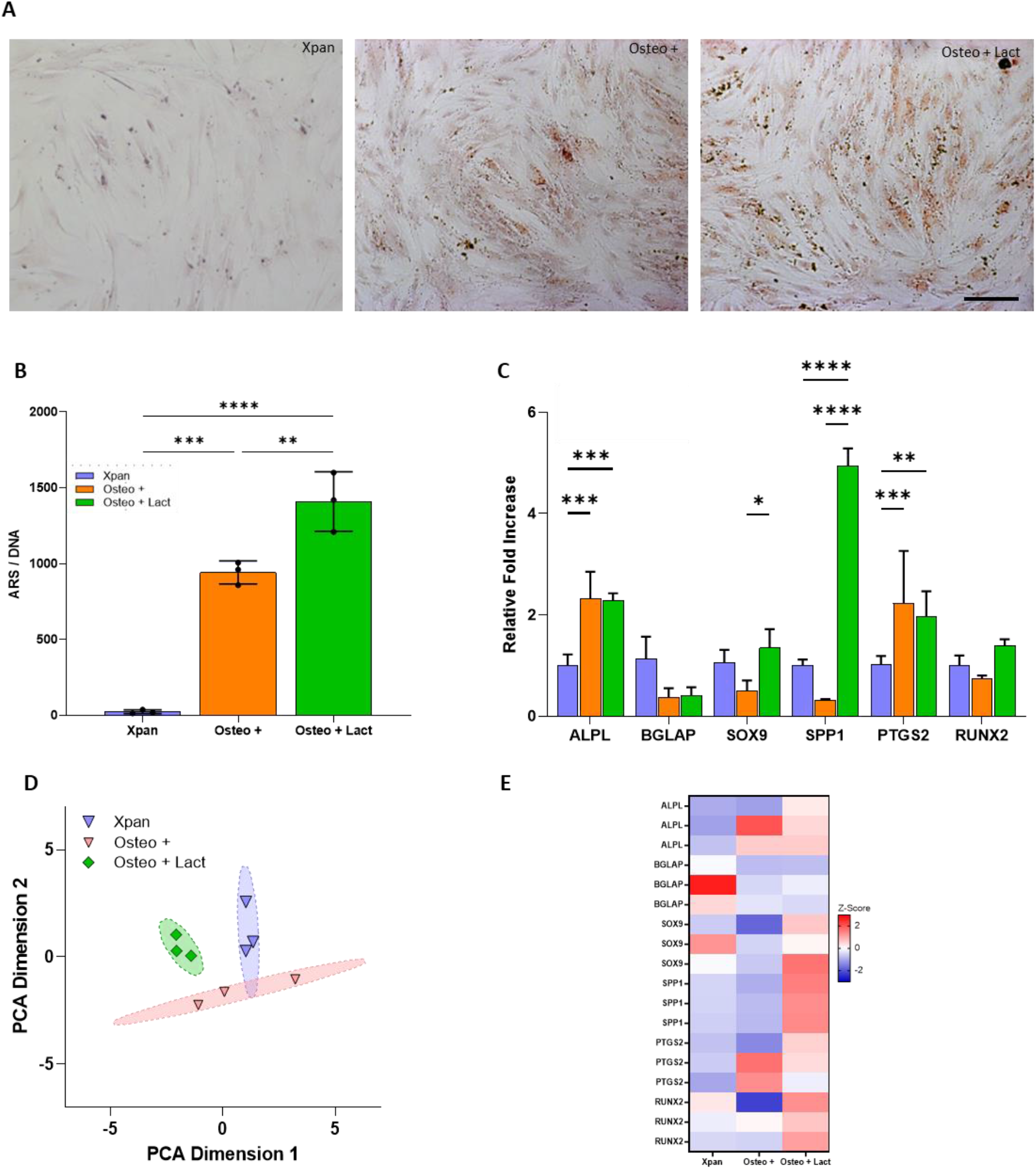
Impact of lactate supplementation of osteogenic differentiation media. **A) Alizarin red staining of hMSCs after 14 days in contact with Osteo +, Osteo + Lact or Xpan medium.** B) Quantification of Alizarin red staining per amount of DNA after 14 days of hMSCs in Xpan, Osteo + and Osteo + Lact conditions. C) hMSCs gene expression of osteogenic markers after 14 days of cell culture. D) PCA distribution of hMSCs osteogenic gene expression after 14 days of cell culture. E) Z-score heatmap of hMSCs gene expression of osteogenic gene markers after 14 days of cell culture. *p-value≤0.05, **p-value≤0.01, ***p-values≤0.001, ****p-value≤0.0001 N≥3

2P-FLIM imaging of NAD(P)H was conducted at day 14 of cell culture (Figure 6A). 2P-FLIM revealed a statistically significant lower τ_avg_ when supplementing osteogenic medium with lactate (1.148 ± 0.005 ns) compared with non-supplemented osteogenic medium (1.187 ± 0.014 ns) (Figure 6B). However, τ_avg_ of Osteo + Lact (1.148 ± 0.005 ns) is statistically significantly higher compared with hMSCs incubated in Xpan medium (1.108 ± 0.002 ns) (Figure 6B). ORR calculations showed statistically significant lower ORR values for Osteo + Lact medium (0.547 ± 0.016 ns) compared with other cell culture medium formulations (0.779 ± 0.016 ns for Xpan and 0.668 ± 0.011 ns for Osteo +) (Figure 6C).

**Figure 6.**
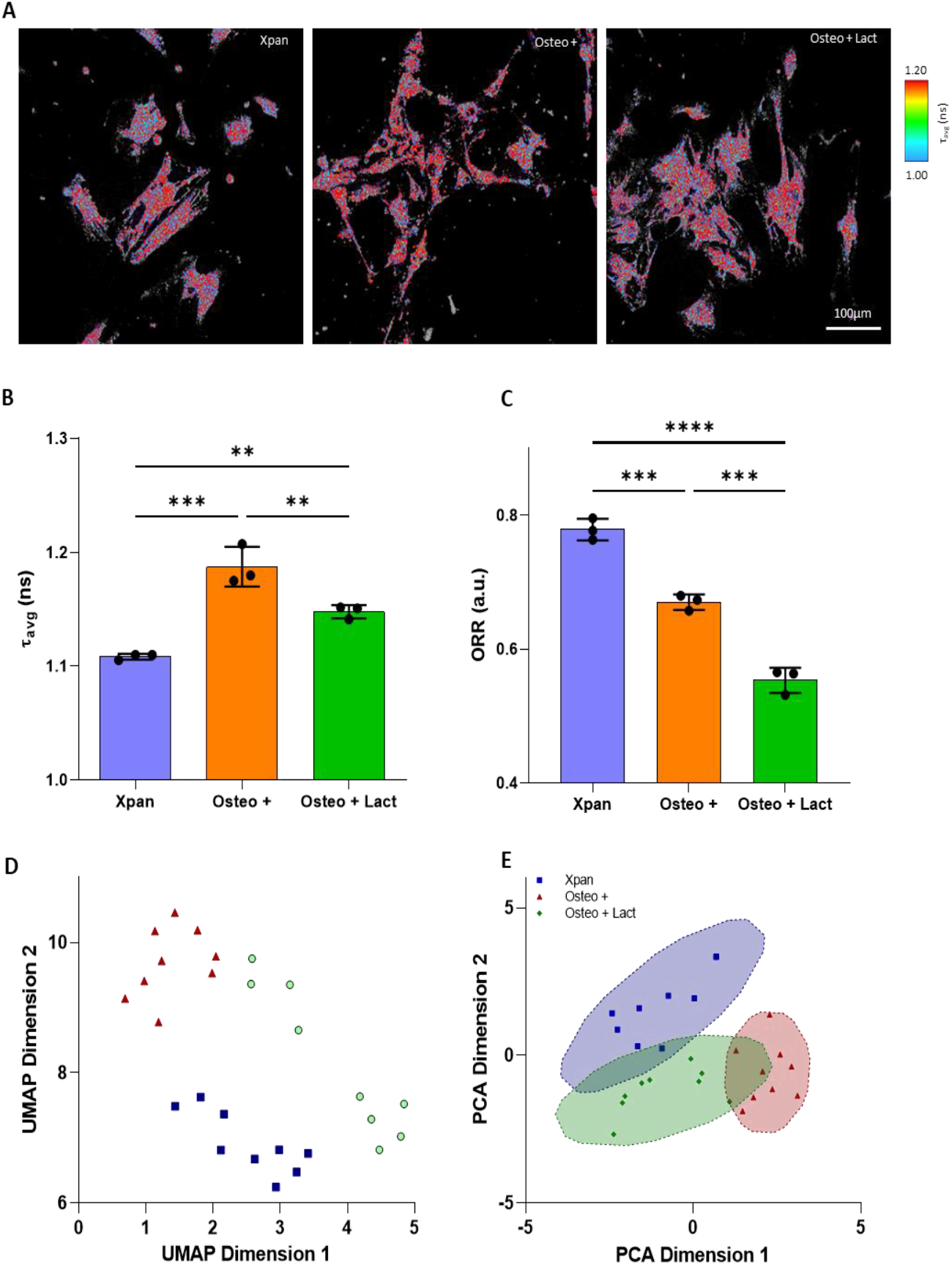
2P-FLIM measurement of lactate supplementation impact on hMSCs metabolism. **A) 2P-FLIM imaging of hMSCs at day 14 of cell culture incubation.** B) Optical Redox Ratio (ORR) of hMSCs at day 0, 3, 7 and 14 of incubation in cell culture medium. C) NAD(P)H protein-bound lifetime τ_1_ at day 14 of cell culture. D) UMAP distribution of 2P-FLIM NAD(P)H and FAD^+^ fluorescence lifetimes and intensity. E) PCA distribution of 2P-FLIM NAD(P)H and FAD^+^ fluorescence lifetimes and intensity. J) hMSCs glucose consumption at day 2, 5, 8 and 11 of cell culture. *p-value≤0.05, **p-value≤0.01, ***p-values≤0.001, ****p-value≤0.0001 N≥3

The decrease in τ_avg_ and decrease of ORR ratios demonstrates a direct metabolic impact on differentiating hMSCs when cell culture medium is supplemented with lactate when compared with non-supplemented osteogenic media. A decrease in τ_avg_ is associated with increased free NAD(P)H and a decrease in ORR is also related with an increase in NAD(P)H fluorescence intensity. The increase in NAD(P)H can be a direct result of increase glutaminolysis resulting in NADH regeneration in the conversion of glutamate to alpha-ketoglutarate as well with NADH production by reconversion of lactate to pyruvate [24, 25].

UMAP distribution analysis of 2P-FLIM results shows a separation between Xpan, Osteo +, Osteo + Lact (Figure 6 D). PCA shows a slight overlap of Osteo + lact with Osteo + and Xpan medium conditions. Nonetheless, statistical analysis of PCA clustering revealed a statistical significant segregation with p-value<0.05 and calculated f-value higher than critical f-value (Figure 6 E, Supplemental table 2).

Regarding metabolite consumption and secretion, lactate supplementation promotes statistically significant lower glucose consumption compared with Xpan medium at early time-points. At day 5 and 8, lactate supplementation of osteogenic medium slightly increases (2.26-fold change) with statistical significance the amount of glucose consumption compared with Osteo + medium (Figure 7 A). At day 11, lactate supplementation promotes a decrease of hMSCs glucose consumption (Figure 7 A). For lactate secretion measurements of lactate supplemented osteogenic medium, exogenous lactate medium contribution was taken in account and the values were corrected. hMSCs incubated in Osteo + Lact conditions have an increasing trend of lactate secretion until day 11 (Figure 7 B). Here, there is a statistically significant higher secretion of lactate when hMSCs are present in Xpan media (202.67 ± 47.84 mM) when compared with the other cell culture conditions (59.33 ± 12.47 mM for Osteo + Lact and 92.67 ± 23.57 mM) (Figure 8B). When hMSCs are incubated in Osteo + Lact medium, there is a higher consumption of glutamine at day 2 (1.827 ± 0.009 mM) compared with other cell culture medium (1.417 ± 0.017 for Osteo + and 1.000 ± 0.041 for Xpan) (Figure 7 C). For the remaining time-points there is a sharp decrease in glutamine consumption for lactate supplemented osteogenic medium (Figure 7 C).

**Figure 7.**
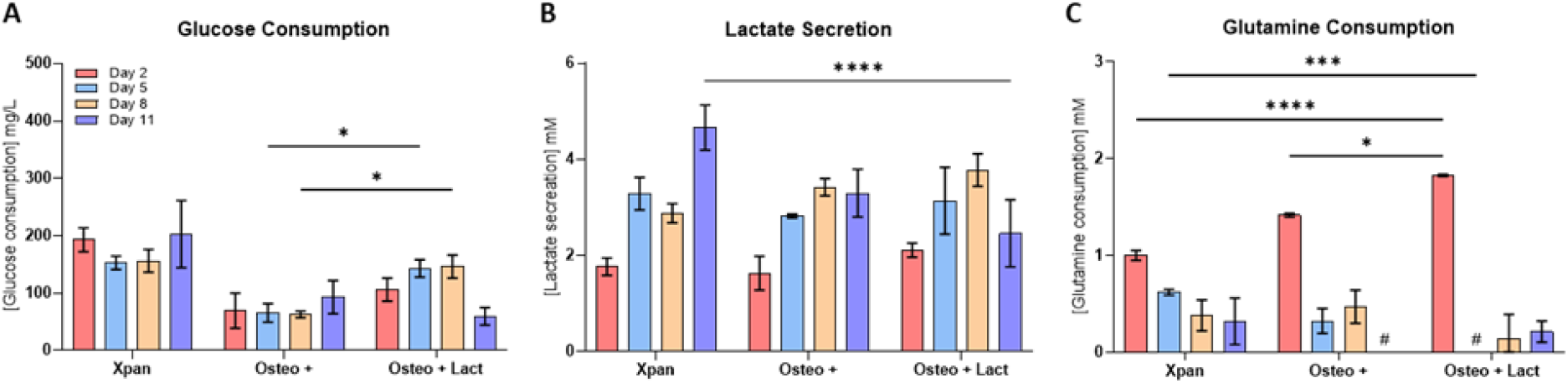
Extracellular metabolite quantification of hMSCs after 14 days in cell culture. **A) hMSCs glucose consumption at day 2, 5, 8 and 11 of cell culture.** B) Lactate secretion at day 2, 5, 8 and 11 of cell culture. C) hMSCs glutamine consumption at day 2, 5, 8 and 11 of cell culture. #no glutamine consumption, *p-value≤0.05, **p-value≤0.01, ***p-values≤0.001, ****p-value≤0.0001 N≥3

**Figure 8.**
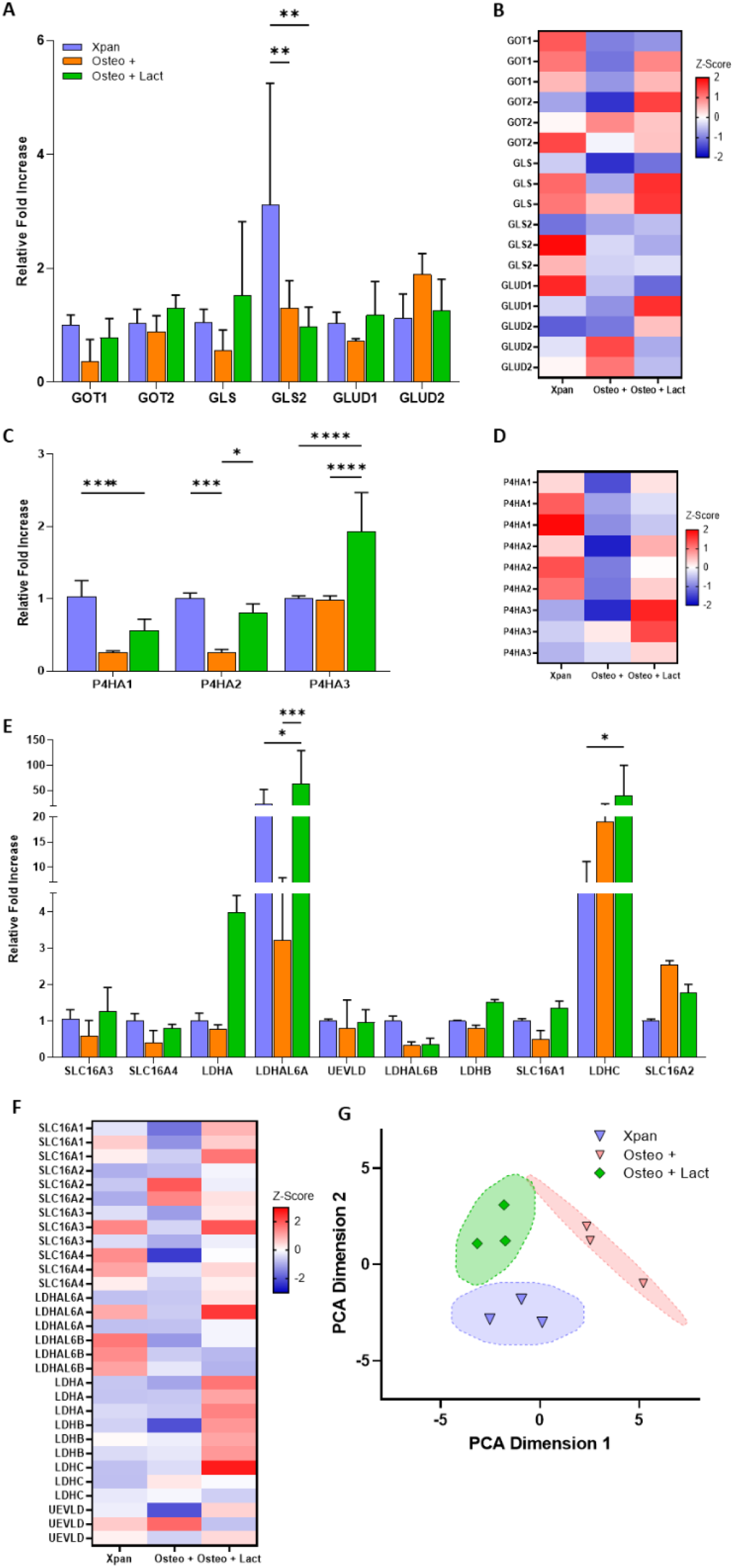
Impact of lactate supplementation of osteogenic differentiation media on the expression of metabolic genes. **A) hMSCs relative fold variation of glutaminolysis genes**. B) Z-score heatmap of genes expression related with glutaminolysis. C) hMSCs relative fold variation of poly-hydrolases genes. D) Z-score heatmap of genes expression of poly-hydrolases. E) hMSCs relative fold variation of lactate transport and metabolism genes. F) Z-score heatmap of genes expression related with lactate transport and metabolism G) PCA distributions of cell culture conditions at day 14 based on glutaminolysis, lactate metabolism and poly-hydrolase gene expression. *p-value≤0.05, **p-value≤0.01, ***p-values≤0.001, ****p-value≤0.0001 N≥3

### Lactate supplementation promotes higher glutaminolysis, lactate metabolism and poly-hydrolases gene expression

Lactate supplementation of osteogenic medium promoted higher yields of osteogenic differentiation validated by osteogenic gene expression and mineral deposition. In addition, lactate supplementation of osteogenic medium promoted lower ORR values and increased glutamine consumption. Here, glutaminolysis, lactate metabolism and poly-hydrolase gene expression was evaluated to establish the impact of lactate supplementation on these metabolic pathways. hMSCs were incubated for 14 days in Xpan, Osteo +, and Osteo + Lact cell culture conditions. Afterwards, the expression of genes related with glutaminolysis, poly-hydrolases, lactate metabolism, lactate transport was measured using qPCR (Figure 8). Osteo + Lact cell culture medium promoted an increase in expression of glutaminolysis related genes such as GOT1 (2.17-fold), GOT2 (1.46-fold), GLS (2.73-fold) and GLUD1 (1.64-fold) when compared with non-supplemented osteogenic medium (Figure 8 A, B). Osteo + Lact induced an increase expression of P4HA1 (2.15-fold), P4HA2 (3.12-fold) and specially P4HA3 (1.97-fold) poly-hydrolases when compared with Osteo + medium (Figure 8 C, D).

Furthermore, Osteo + Lact medium also promoted the increase expression of lactate transporter genes such as SLC16A1 (2.78-fold), SLC16A3 (2.15-fold), SLC16A4 (2.00-fold) and lactate metabolic genes LDHAL6A (19.89-fold), LDHAL6B (1.09-fold), LDHA (5.16-fold), LDHB (1.89-fold), LDHC (2.12-fold) (Figure 8 E, F). Finally, PCA distribution of cell culture conditions calculated based on the expression of metabolism related genes showed a statistical significant segregation between Osteo +, Osteo + Lact and Xpan cell culture conditions with p-value<0.05 and calculated F-value higher than critical F-value (Figure 8 G).

### Blocking glutaminolysis and lactate absorption impacts osteogenic differentiation

Lactate supplementation of osteogenic medium promoted higher hMSCs mineral deposition and upregulation of glutaminolysis and the reconversion from lactate to pyruvate as measured by τ_avg_, ORR, metabolites consumption and gene expression. To validate that osteogenic differentiation is dependent on glutaminolysis and lactate transport/metabolism, these metabolic pathways were blocked using chemical inhibitors. To inhibit glutaminolysis BPTES; a chemical competitor of glutamate synthase enzyme, the first step of the glutaminolysis pathway was used. Inhibiting this enzyme has showed to inhibit the glutaminolysis pathway [26]. Regarding lactate transport, α-CHC was used. This chemical component inhibits the uptake of lactate by blocking SLC16A1 transporter [27]. This transmembrane transporter was shown to be the main entry point of lactate in hMSCs cells [28] (Figure 9 A).

**Figure 9.**
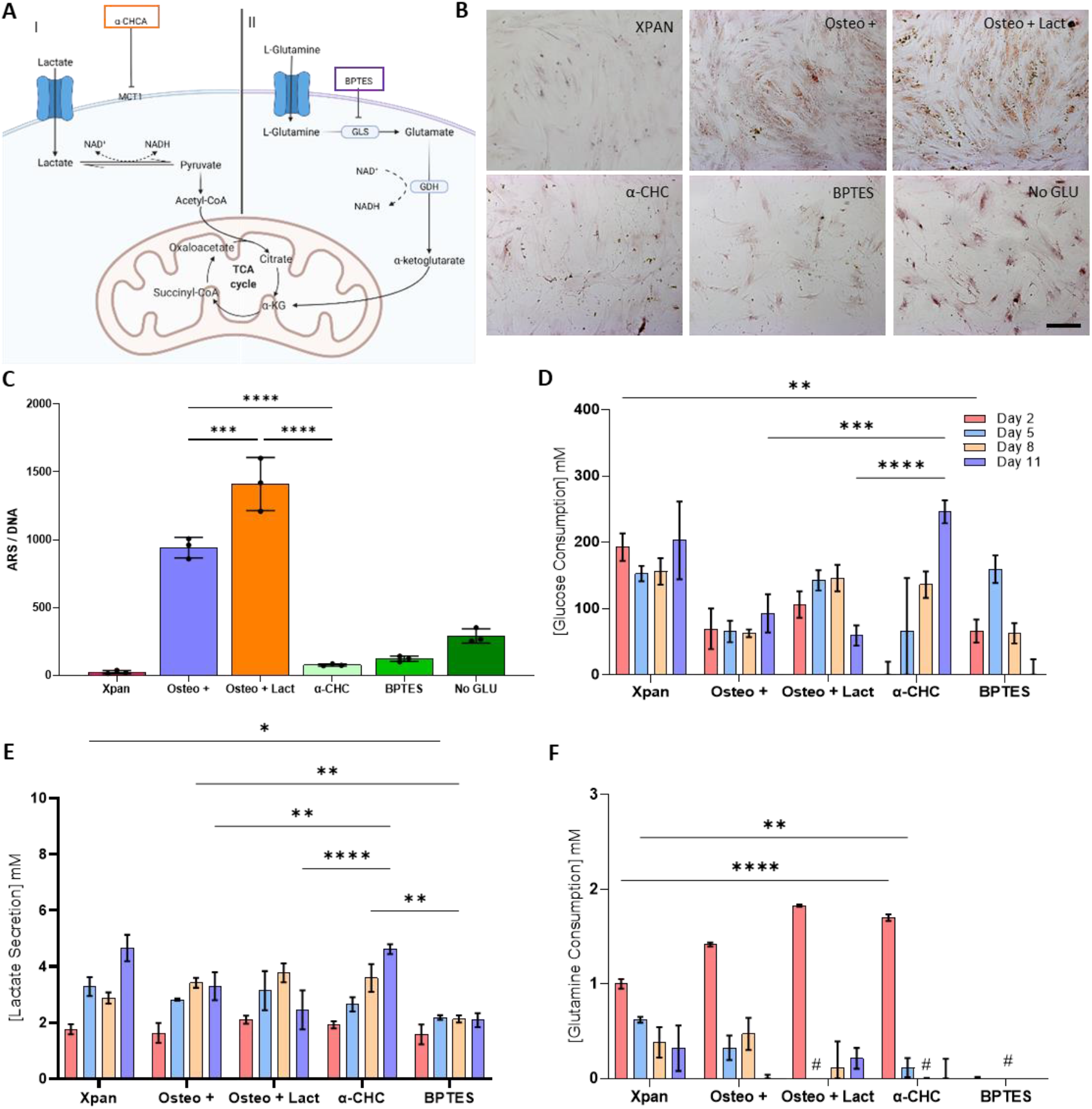
Inhibition of osteogenic differentiation using osteogenic medium supplemented with BPTES or α-CHC during 14 days of cell culture. A) Schematic detailing the inhibition target. B) Alizarin red staining (ARS) of hMSCs after 14 days in contact with Osteo +, Osteo + Lact, α-CHC or BPTES supplemented osteogenic medium, Osteo + medium without glutamine (No Glu) or Xpan medium. C) Quantification of Alizarin red staining per amount of DNA after 14 days of hMSCs in various cell culture conditions. D) hMSCs glucose consumption at day 2, 5, 8 and 11 of cell culture. E) Lactate secretion at day 2, 5, 8 and 11 of cell culture. F) hMSCs glutamine consumption at day 2, 5, 8 and 11 of cell culture. #no glutamine consumption, *p-value≤0.05, **p-value≤0.01, ***p-values≤0.001, ****p-value≤0.0001 N≥3

When treating hMSCs during 14 days with Osteo + medium supplemented with α-CHC, BPTES or no glutamine added (No Glu), there is a reduction of alizarin red staining demonstrating lower mineral deposition (Figure 9 B). We further validated this result by calculating the amount of alizarin red stain per amount of DNA. Here, we noticed a statistically significant decrease of alizarin red staining when hMSCs are incubated in Osteo + medium supplemented with either α-CHC or BPTES (Figure 9 C).

α-CHC treatment promotes an increasing trend of glucose consumption being statistically significantly higher at day 11 when compared with Osteo + medium and Lactate supplemented Osteo + medium. BPTES has an increasing trend of glucose consumption until day 5. Afterwards, there is a decrease of glucose consumption until day 11 (Figure 9D). α-CHC treated hMSCs have an increasing trend of lactate secretion during the period of incubation. At day 11, there is a statistically significant higher lactate secretion when compared with Osteo + or Lactate supplemented Osteo + medium. For BPTES, there is a stable lactate secretion during the incubation period. In addition, BPTES treatment promotes a statistically significant decrease of lactate when compared with all of the other cell culture medium conditions (Figure 9E).

α-CHC supplemented osteogenic medium promotes a decreasing trend of glutamine consumption whilst being statistically significantly higher when compared with Xpan at day 2. At later time-points, a-CHC treated hMSCs have a decreasing trend of glutamine consumption (Figure 9F). As expected, BPTES supplementation of osteogenic medium has a negative impact on glutamine consumption. With BPTES treatment hMSCs did not consume any glutamine during cell culture (Figure 9F).

2P-FLIM of NAD(P)H was performed to assess the impact of α-CHC and BPTES on hMSCs metabolism after 24 hours treatment (Figure 10 A). hMSCs τ_avg_ and ORR measurements are not directly impacted by α-CHC treatment, resulting in no statistical significance in all cell culture conditions for both τ_avg_ and ORR parameters (Figure 10 B, C). For BPTES, the presence or removal of glutamine resulted in distinct metabolic profiles and impacted τ_avg_ and ORR measurements. The removal of glutamine from osteogenic medium statistically significantly decreased τ_avg_ (0.90-fold decrease) of hMSCs compared with Xpan, Osteo+ and Osteo+ Lact cell culture medium condition (Figure 10 D). Regarding ORR, BPTES treatment of osteogenic medium and the removal of glutamine from osteogenic medium resulted in an increase of 1.10-fold and 1.17-fold of ORR, respectively, when compared with other cell culture medium conditions (Figure 10 E).

**Figure 10.**
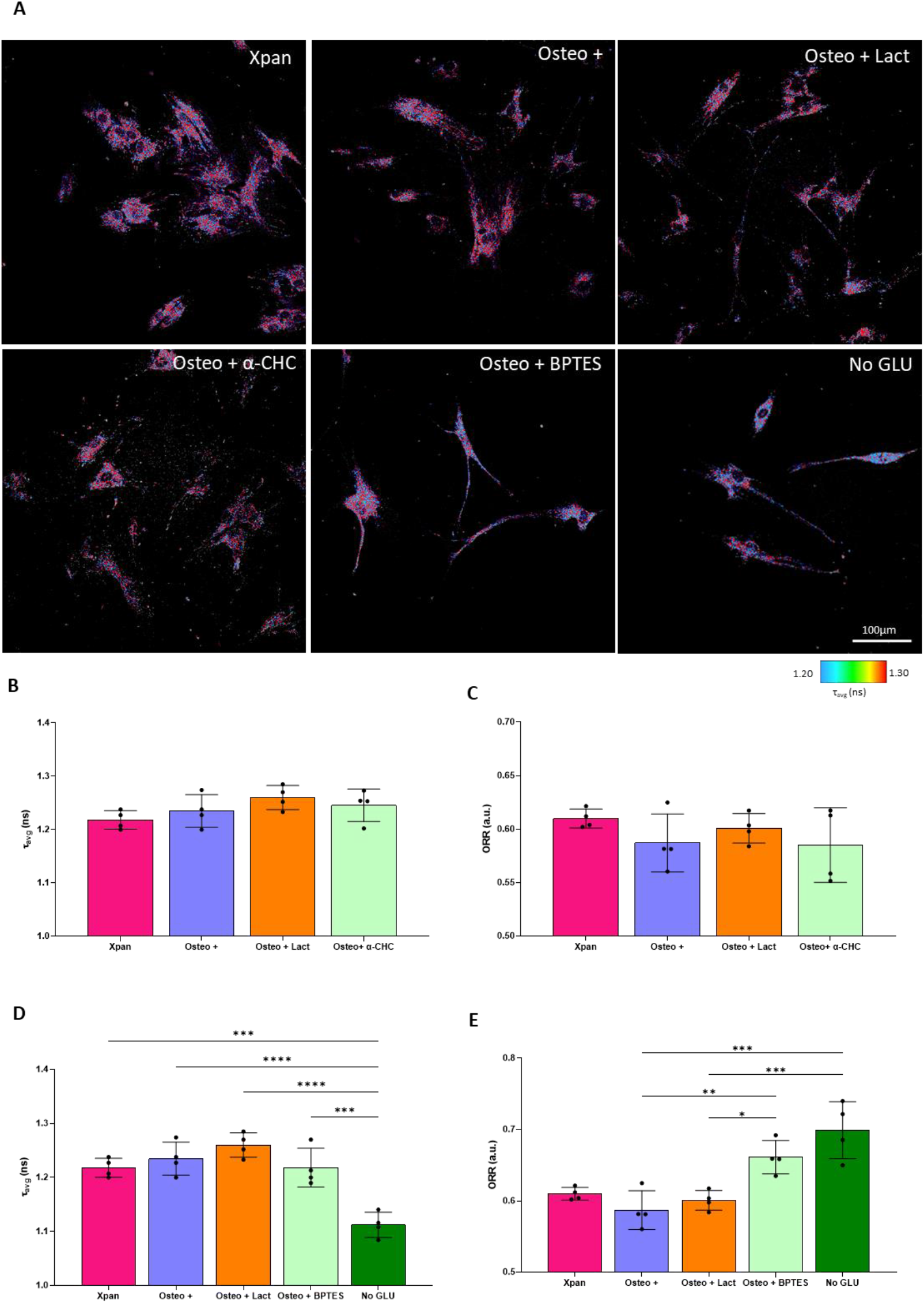
2P-FLIM validation of metabolic inhibition by α-CHC and BPTES after 24 hours. A) 2P-FLIM imaging of hMSCs in different cell culture conditions B) NAD(P)H τ_avg_ measurement of hMSCs after 24 hours of α-CHC treatment. C) Optical Redox Ratio (ORR) of hMSCs after 24 hours of α-CHC treatment. D) NAD(P)H τ_avg_ measurement of hMSCs after 24 hours of BPTES treatment. E) Optical Redox Ratio (ORR) of hMSCs after 24 hours of BPTES treatment. *p-value≤0.05, **p-value≤0.01, ***p-values≤0.001, ****p-value≤0.0001 N≥3

## Discussion

In this study, hMSCs osteogenic differentiation was achieved using osteogenic medium and validated by mineral deposition and osteogenic gene expression. During the osteogenic differentiation, 2P-FLIM was used to monitor the metabolic profile of hMSCs. This non-invasive technique showed that hMSCs in osteogenic medium are more dependent on OxPhos as a source of energy than hMSCs in Xpan medium. Also, 2P-FLIM highlighted a higher dependence on glutaminolysis and anabolic pathways verified by increased glutamine consumption. Osteogenic medium was supplemented with lactate to upregulate glutaminolysis and other anabolic pathways. Here, lactate supplementation of osteogenic medium drove increased mineral deposition and higher osteogenic gene expression when compared with non-supplemented osteogenic medium. Again, 2P-FLIM revealed a higher dependence of anabolic pathways and higher ratio of free NAD(P)H. The higher ratio of free NAD(P)H is a consequence of glutaminolysis and re-conversion of lactate into pyruvate. Osteogenic differentiation dependence on glutaminolysis and lactate re-conversion to pyruvate was validated by inhibiting glutaminolysis using BPTES and lactate uptake using α-CHC. The use of each metabolic inhibitor resulted in a decrease in mineral deposition and metabolite uptake. Furthermore, it was demonstrated that inhibiting glutaminolysis resulted in higher ORR ratios, validating that lower ORR ratios can result from higher glutaminolysis rates.

2P-FLIM of hMSCs osteogenic differentiation uncovered a metabolic trends towards OxPhos and a higher dependence on anabolic pathways. hMSCs treated with osteogenic medium showed a higher τ_avg_, as early as day 3, reflective of a metabolic shift towards OxPhos as a consequence of a higher proportion of longer lifetime and protein-bound NAD(P)H [9, 29] (Figure 2F). Also, τ_1_ lifetimes are higher in osteogenic conditions, highlighting higher amounts of NADPH and anabolic activity [10, 30] (Figure 2G). Varying interpretations of ORR are reported in previous studies [31, 32] and for this study, we adopt a lower ORR reflecting a higher fraction of NAD(P)H and a lower FAD^+^ associated with upregulation of Krebs cycle. An increased generation of NADH will result in a decreased ORR (Equation 3)[33, 34]. Osteogenic medium induced lower ORR possibly to fuel both OxPhos and anabolic pathways (Figure 2J). These metabolic results are in agreement with the literature; higher OxPhos and anabolic pathways dependence during osteogenic differentiation [5, 19, 35]. However, we are the first to uncover a metabolic switch as early as day 3. UMAP and PCA distribution analysis of 2P-FLIM variables confirmed a time-dependent metabolic switch during osteogenic differentiation. Taking into account 2P-FLIM variables in the UMAP analysis, there is noticeable a separation between Xpan and Osteo + treated cells at day 7 and day 14. PCA analysis revealed also a separation at day 7 and 14 between these culture conditions. Overall, the statistical significance and the distance between PCA and UMAP clusters of both cell culture conditions increases with longer incubation periods. Mahalanobis distance, Two-sample Hotellings T2 test, F-value, F-critical and P-values were calculated to ensure statistical significance between PCA cluster groups (Supplemental table 2). The clear and statistically valid segregation of cell culture and hMSCs profiles demonstrate the possibility to use PCA and UMAP not only as data-visualisation tools but to possibly be used as classification tools. The trends observed with 2P-FLIM and extracellular metabolite quantification, confirm a metabolic switch towards OxPhos by reduction of glucose consumption while showing a higher dependence of glutaminolysis at earlier time-points. The metabolic switch towards a more OxPhos dependence is a known feature of differentiated cells and it has been demonstrated to occur also during osteogenic differentiation [5, 36]. Interestingly, glutamine has been shown to be a critical metabolite to achieve osteogenic differentiation [8]. One study by Yu et al., demonstrated that glutamine is an important metabolite to achieve osteogenic differentiation by its anaplerotic role in feeding the Krebs cycle by proxy of alpha-ketoglutarate. In addition, this study demonstrated that glutamine removal impacts osteogenic differentiation by decrease of alizarin red staining [8]. Interestingly, Gayatri et al., demonstrated that increasing glutamine concentrations 10-fold supresses osteogenic differentiation of human and murine MSCs [37]. Therefore, to promote glutaminolysis and anaplerotic pathways and consequently promote earlier osteogenesis, the glutaminolysis pathway rate needs to be increased without increasing extracellular concentrations of glutamine.

To achieve a higher rate of glutaminolysis, we decided to upregulate the glutaminolysis pathway by modifying the Osteo + cell culture medium formulation. Pérez-Escuredo et al., showcase the upregulation of glutamine uptake and glutaminolysis promoted by increased lactate concentration on oxidative tumour cells. Pérez-Escuredo et al., study demonstrates how extracellular lactate can promote increased glutaminolysis [23]. Following this approach, we decided to increase exogenous lactate levels in the Osteo + medium to promote higher glutaminolysis and hMSCs osteogenic differentiation and mineral deposition. To account for pH changes due to increased lactate concentration, exogenous lactate in the form of sodium lactate was used to stabilize pH values [23]. Lactate supplementation of osteogenic media yielded a higher mineral deposition and osteogenic differentiation (Figure 3A, B, C, D). Here, increased gene expression of ALPL, SOX9, SPP1 and RUNX2 are upregulated markers of hMSCs osteogenesis and follow the trend observed in literature [38, 39]. Using 2P-FLIM, we observed a shift in metabolism when supplementing osteogenic medium with lactate (Figure 6A). A lower τ_avg_ shows an increase in free NAD(P)H fraction in the cytoplasm concomitant with a reduction of protein bound-NADH. This shift in NAD(P) fraction is associated with higher metabolic dependence on glycolysis (Figure 6 B). Simultaneously, there is a decrease in ORR associated with OxPhos and anaplerotic metabolic pathways such as glutaminolysis (Figure 6 C). UMAP and PCA distribution analysis of 2P-FLIM data revealed a segregation between the hMSCs treated with Osteo + SL, Osteo + and Xpan medium (Figure 6 D, E). This shows that the three cell culture formulations promote distinct metabolic properties in hMSCs. Metabolic gene expression revealed for most genes, higher expression levels when hMSCs are incubated for 14 days in osteogenic lactate supplemented cell culture medium (Figure 8). These glutaminolysis genes are responsible for the conversion of glutamine to glutamate and the conversion of glutamate to alpha-ketoglutarate in the cytoplasm and the mitochondrial matrix [40, 41]. These poly-hydrolases are located within the endoplasmic reticulum, are activated by the presence of alpha-ketoglutarate and are crucial for the formation of the collagen triple helix. The incomplete hydroxylation of proline residues impacts collagen triple helix formation reducing collagen secretion to the cytoplasm and lastly reducing extracellular secretion of collagen [42]. The ability to secrete high amounts of extracellular matrix such as collagen I is a known feature of osteoblasts and important during bone formation [43]. Lactate metabolic genes are also highly expressed in lactate supplemented osteogenic medium when compared with non-supplemented osteogenic medium. SL16A1 in particular has been showed to express MCT1 one of the main membrane transporter of lactate in hMSCs [44]. LDH is also highly expressed in all isoforms when treating hMSCs with Osteo+ cell culture medium. LDH is responsible for conversion of pyruvate to lactate and is capable of performing the reverse enzymatic reaction [45]. The upregulation of LDH gene expression without an increase in extracellular lactate concentrations demonstrates that LDH has a higher ratio of reverse enzymatic reaction compared with the forward reaction. Glutaminolysis and lactate gene expression demonstrate that exogenous lactate supplementation of lactate medium promotes upregulation of glutaminolysis while increasing lactate uptake and re-conversion to pyruvate in hMSCs. PCA metabolic gene expression analysis revealed a statistically significant segregation between all cell culture conditions (Figure 8 G). This confirms that lactate supplementation of osteogenic medium is having a direct impact on hMSCs metabolism. hMSCs when treated with Osteo + Lact medium have higher level of glutaminolysis rate, higher uptake of lactate, higher rates of lactate conversion to pyruvate and higher secretion levels of collagen. These metabolic shifts can be therefore responsible for higher levels of osteogenic differentiation.

After showing the importance of glutaminolysis and lactate in osteogenic differentiation we decided to block glutaminolysis using BPTES or removing glutamine (No Glu) from the osteogenic medium, as well as blocking the uptake of lactate using α-CHC (Figure 9 A). BPTES impacts directly the GLS enzyme whilst α-CHC inhibits the SLC16A1 transporter, responsible for lactate uptake. Alizarin red staining and extracellular metabolic analysis shows that inhibiting lactate uptake and glutaminolysis impacts negatively osteogenic differentiation. 2P-FLIM NAD(P) measurements of the metabolic profile of hMSCs after 24 hours treatment demonstrated that α-CHC does not impact either τ_avg_ or ORR redox ratio at earlier treatments times. Since α-CHC directly inhibits a transmembrane transporter responsible for lactate uptake, the impact on metabolism might not be pronounced in short culture times probably due to the pyruvate available on the medium which can evade lactate uptake to be converted back to pyruvate while generating NADH. However, BPTES treatment or removal of glutamine significantly impacted hMSCs metabolism after 24 hours. Removal of glutamine produced a statistically significant decrease of τ_avg_ possibly generated by the cell compensating the absence of an essential metabolite by increasing glycolysis to fuel the Krebs cycle for cellular differentiation. ORR measurements show a statically significant increase in BPTES and no glutamine osteogenic medium. This trend is a direct opposite from what is observed experimentally with upregulated glutaminolysis in this study. Increased ORR due to BPTES or removal of glutamine results from a reduction of NADH availability due to not being regenerated during glutamate conversion to alpha-ketoglutarate [31]. Lactate was added to osteogenic medium in the presence of BPTES and α-CHC in order to complement glutaminolysis and lactate transport inhibition studies in hMSCs. Lactate supplementation of osteogenic medium with BPTES was not able to rescue osteogenic differentiation. In addition, lactate supplementation of osteogenic medium with α-CHC was also not able to promote osteogenic differentiation (Supplemental figure 2). This results demonstrates that osteogenic differentiation supplemented with lactate impacts directly both metabolic pathways: glutaminolysis and lactate uptake.

The culmination of these results validate the role of glutaminolysis in hMSCs osteogenic differentiation following the trend observed by Yu et al. in which blocking glutaminolysis with BPTES negatively impacted osteogenic differentiation [8]. It seems that both metabolic pathways are essential for osteogenic differentiation, glutaminolysis metabolic pathway and the direct and indirect role of lactate in hMSCs metabolism. In this study, a new role for lactate in osteogenic differentiation of hMSCs was uncovered. Here, lactate can act indirectly on osteogenic differentiation as a signalling to upregulate glutaminolysis and directly by being used as metabolic fuel to regenerate pyruvate. In particular, this study demonstrates for the first time the importance of lactate for hMSCs differentiation. Only recently, has lactate started to be understood as a fuel source and a distinct signalling molecule instead of just a metabolic by-product. Lactate is becoming an interesting metabolite and has been showed to contribute to reduce inflammatory cell signalling [46]. Furthermore, the only study demonstrating the impact of extracellular lactate on stromal cells was a study by Gatie et al. In this study, Gatie et al., observed that increasing extracellular lactate stimulates murine embryonic stromal cells differentiation towards extraembryonic endoderm cells *in vitro* [47].

This study demonstrates the possibility to modulate hMSCs differentiation using extracellular metabolites. The importance of glutaminolysis in osteogenic differentiation was verified. As well, new roles for lactate metabolite were uncovered while promoting increased hMSCs osteogenic differentiation. This work demonstrates not only the importance of essential metabolites in osteogenic differentiation but also showcases 2P-FLIM as a method to uncover and validate metabolic profiles of hMSCs non-invasively in real-time. This new metabolic viewpoint in stromal cell differentiation should be taken in account when promoting stromal cell differentiation in vitro. Importantly, the metabolic environment during cell culture or cell maturation should be established to either promote higher differentiation rates of hMSCs or to simulate metabolic properties of native tissue.

In conclusion, we show that osteogenic differentiation is a phenotypic and metabolic process that occurs in a time-dependent manner reliant on glutaminolsysis and lactate. The inhibition of these pathways using BPTES and α-CHC and withdrawal of glutamine negatively impacts osteogenic differentiation. The supplementation of osteogenic medium with lactate promoted higher osteogenic differentiation, glutaminolysis, lactate metabolism and end-point poly-hydrolises gene expression. At later time-points, lactate supplemented osteogenic cells exhibit distinct metabolic profiles when compared with cells in non-supplemented osteogenic medium and Xpan growth medium. This study provides information on the metabolic shifts occurring during hMSCs differentiation as well the role of glutaminolysis and extracellular lactate in osteogenic differentiation. Potentially, the use of lactate as a coating or as a biocompatible material in the form of lactic acid can promote further stromal cell differentiation at an implantation site. The metabolic role of lactate as fuel and signalling molecule is still a new approach to a metabolite that is traditionally considered as a metabolic by-product and established as a cellular metabolism endpoint. Current research is now shining a new light on the impact of exogenous lactate impact on cellular function. However, research in various cell types using numerous methodologies is still require to establish the various roles and potential impact of lactate.

## Experimental Procedures

### Cell Culture

Human mesenchymal stromal cells (hMSCs) were purchased from Lonza^®^ were cultured at passage 3-4 in expansion media (Xpan) prepared with Dilbecco’s Modifed Eagle’s Medium (DMEM), 1000 mg/L low glucose (Sigma-Aldrich) containing 10% fetal bovine serum (FBS) (Gibco^®^ by Life Technologies) and 2% Penicilin Streptomycin (Pen-Strep) (Sigma Aldrich) at 37°C with 5% CO2. To initiate osteogenic differentiation, osteogenic media (Osteo +) was prepared by adding 0.2% (v/v) dexamethasone, 1% (v/v) β-glycerol phosphate and 0.029% (v/v) ascorbic acid as supplements to Xpan media.

For lactate supplementation of Osteo + media, 75 mM of lactate was used as a sodium salt, to avoid changes in extracellular pH (Osteo + Lact). For metabolic inhibition studies, either 5 mM of α-CHC or 10 μM BPTES were added to Osteo + media.

### Two-photon fluorescence lifetime imaging microscopy (2P-FLIM)

2P-FLIM was performed on hMSCs (at passage 3-4) seeded in 35 mm petri dishes. 2P-FLIM was achieved using a custom upright (Olympus BX61WI) laser multiphoton microscopy system equipped with a pulsed (80 MHz) titanium:sapphire laser (Chameleon Ultra, Coherent^®^, USA), water-immersion 25× objective (Olympus, 1.05NA) and temperature-controlled stage at 37 °C. Two photon excitation of NAD(P)H and FAD^+^ fluorescence was performed at the excitation wavelength of 760 and 800 nm, respectively. A 458/64 nm and 520/35 nm bandpass filter were used to isolate the NAD(P)H and FAD^+^ fluorescence emissions based on their emission spectrum.

512 × 512 pixel images were acquired with a pixel dwell time of 3.81 μs and 30 second collection time. A PicoHarp 300 TCSPC system operating in the time-tagged mode coupled with a photomultiplier detector assembly (PMA) hybrid detector (PicoQuanT GmbH, Germany) was used for fluorescence decay measurements, yielding 256 time bins per pixel. TCSPC requires a defined “start”, provided by the electronics steering the laser pulse or a photodiode, and a defined “stop” signal, realized by detection with single-photon sensitive detectors. The measuring of this time delay is repeated many times to account for the statistical variance of the fluorophore’s emission. For more detailed information, the reader is referred elsewhere [48]. Fluorescence lifetime images with their associated decay curves for NAD(P)H were obtained, with a minimum of 1×10^6^ photons peak, and region of interest (ROI) analysis of the total cells present on the image was performed in order to remove any background artefact. The decay curved was generated and fitted with a double-exponential decay without including the instrument response function (IRF) (Equation (1)).

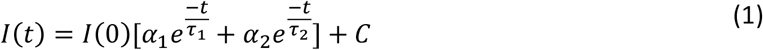

I(t) represents the fluorescence intensity measured at time t after laser excitation; α1 and α2 represent the fraction of the overall signal proportion of a short and long component lifetime, respectively. τ_1_ and τ_2_ are the long and short lifetime components, respectively; C corresponds to background light. Chi-square statistical test was used to evaluate the goodness of multi-exponential fit to the raw fluorescence decay data. In this study, all of the fluorescence lifetime fitting values with χ^2^ < 1.3 were considered as ‘good’ fits. For NAD(P)H, the double exponential decay was used to differentiate between the protein-bound (τ_1_) and free (τ_2_) NAD(P)H. The average fluorescence lifetime was calculated using Equation (2).

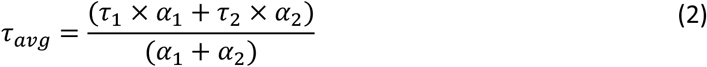

Intensity-based images of NAD(P)H and FAD^+^ were acquired, and their ratio was calculated using Equation (3) to obtain the optical redox ratio (ORR).

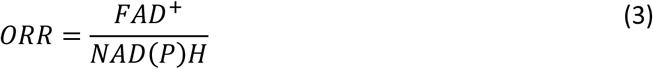

### qPCR analysis

hMSCs were lysed and re-suspended in 1 ml of TRIzol, briefly vortexed and 200 μL of chloroform was added. Samples were centrifuged at 12,000g for 15 minutes at 4°C to achieve phase separation. The aqueous phase was separated to a fresh RNAse free microtube. RNA precipitation was performed by adding 2 μL of glycol-blue and 500 μL of isopropanol. RNA pellets were washed using 1 mL of 70% of ethanol. RNA pellets were re-suspended in 20 μL of RNAse free water. RNA yield and purity was measured using a NanoDrop^™^. To estimate total RNA concentration, absorbance values obtained at 260 nm were obtained. 260/280 and 260/230 absorbance ratios were calculated to estimate RNA purity. A 260/280 ratio of *1.8 and a 260/230 ratio of *1.8 were deemed acceptable for RNA purity. cDNA transcription was performed with High capacity cDNA reverse transcription kit from Applied Biosystems^™^ according to manufactures instructions. A master mix solution was prepared by adding RNase inhibithor, Multiscribe^™^ reverse transcriptase enzyme, RT^®^ buffer, random primers and 100 mM of dNTP Mix. RNA samples were added to the mastermix to achieve a reaction volume of 20 μL. A thermal cycle was performed with a heating step of 25 °C for 10 minutes, 37 °C for 120 min, followed by 85 °C for 5 minutes. A 100% conversion rate was assumed to determine cDNA concentration.

Gene expression was analysed using a custom PrimePCR^®^ designed plate using SYBR^®^ Green primers according to manufactures guidelines. Here, 10 μL of Universal SYBR^®^ Green Supermix is added to each well of the 96-well plate to reconstitute the dried PCR primers. 4 μL of cDNA samples were added to each well and nuclease-free water was used to achieve 20 μL of reaction volume. All primers were purchased from Bio-Rad laboratories. Accession numbers, Unique Assay ID, Amplicon Length, Gene name and symbol are present in Supplemental Table 1. Primer sequences remain Bio-Rad laboratories proprietary information and are not available. PCR primers specificity are verified and guaranteed by the manufacturer. qPCR was performed on a fastPCR 96-well plate instrument (Applied Biosystems) to obtain comparative ΔΔCt values. Gene expression was compared between samples using the 2^-ΔΔCt^ method [49]. QPCR data was normalized to β-actin gene expression.

Z-score heatmaps of gene expression values were generated by calculating the z-score of each sample and plotted in a heatmap. Z-score values are obtained using equation 4:

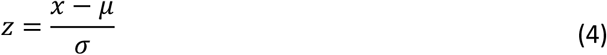

The z-score (z) is calculated using *x* the observed value, μ the mean value of the sample and σ is the standard deviation of the sample

### Metabolite Analysis

#### Glucose

For metabolite analysis, cell culture medium was collected at 2, 5, 8 and 11 days after being in contact with cell cultures. For extracellular measurement of glucose, the ChromaDazzle^™^ Glucose Assay kit from AssayGenie was used. Here, glucose standards were prepared by dissolving glucose stock solution 300 mg/dL in distilled water with concentrations ranging from 0 to 300 mg/dL to prepare a standard curve. Cell culture medium samples and standards were diluted 1/100 in o-toluidine reagent from AssayGenie. Samples were incubated in a boiling water bath for 8 minutes and cooled down in ice for 4 minutes. 200 μL were added to clear-bottom 96 well plates and absorbance values were measured in a microplate reader at 630 nm.

#### Lactate

Extracellular measurements of lactate was performed using an L-Lactic acid colorimetric assay by AssayGenie. Here, lactate standards were obtained by sequentially diluting a 10 mM/L lacatate standard to obtain a standard curve with the concentration ranging from 0 to 7 mM/L. Samples and standards were added directly to 96 well plates and diluted 60x in chromogenic solution. For samples with increased exogenous lactate, 12x dilutions were performed. Samples were incubated at 37 °C for 5 minutes and read in a microplate reader at 530 nm.

#### Glutamine

Extracellular measurements of glutamine were performed using ChromaDazzle^™^ Glutamine Assay kit from AssayGenie. This kit is based on the hydrolysis of glutamine to glutamate and formation of a colorimetric product. First, a standard curve was calculated by diluting glutamine stock solution 100 mM in distilled water. Standards with concentrations from 0 to 2 mM were obtained. Enzymes A qnd B from the kit were reconstituted according to manufacturer’s instructions. Extracellular culture medium and standards were added to clear-bottom 96-well plates and diluted 10x in assay solution containing enzymes. The enzymatic reaction was stopped at 40 min at room temperature with a stop reagent and absorbance values were measured in a plate reader at 550 nm.

In this study, there is an interest in measuring glucose and glutamine consumption. Therefore, to calculate metabolite concentration equation 5 was used. For glucose initial concentration was assumed 1000 mg/L and 2.0mM for glutamine concentration according to manufactures formulations. Extracellular lactate concentration measurements are presented directly without using equation 4. In addition, for lactate measurements in lactate supplemented osteogenic medium, were corrected by taking in account the amount of lactate added exogenously.

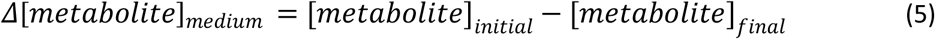

### PCA and UMAP

Uniform Manifold Approximate and Projection (UMAP) and Principal Components Analysis (PCA) were used as data visualisation tools to visualize clustering within 2P-FLIM data variables (Python). For this analysis, all images acquired (at least 3 per sample and condition) were used as plot points and 2P-FLIM variables or gene expression data were used as variables. Covariance error ellipses with confidence values of 99% were calculated and plotted in PCA graphs. Mahalanobis distance, two-sample Hotellings T^2^ statistic, F-value, Critical F-Value and P-values were calculated to ensure statistical significant separation of PCA clusters [50]. (Supplemental Table 2)

### Alizarin red staining (ARS)

hMSCs were fixed with 4% paraformaldehyde (PFA) and alizarin red staining was performed for 30 minutes with a solution of 1% alizarin red in distilled water and imaged with a 10x objective using an Olympus BX41TF brightfield microscope. Alizarin red molecules bind to calcium molecules and generate a birefringent insoluble red complex [51]. Alizarin red stained cells were dissolved for 30 minutes in 10 % (v/v) acetic acid. Cells were lifted with a cell scrapper and transferred to microtubes and heated until 85 °C for 10 minutes. The tubes were centrifuge at 20,000 g for 15 minutes and 10% (v/v) ammonium hydroxide was added. Samples were pipetted in triplicate on a 96 well plate and absorbance was measured in a plate reader at 405nm.

### DNA quantification

For sample lysis, hMSCs were rinsed twice using cold phosphate buffer saline solution (PBS) and were digested using a lysis buffer containing 0.2% v/v Triton X100, 10 mM Tris pH 8, 1 mM EDTA and DNAse free water. Afterwards, the cells suspended in lysis buffer were transferred to microtubes and sonicated for 60 seconds to ensure complete cellular lysis.

DNA quantification was performed using a Picogreen^™^ biochemical assay on digested samples according to manufacturer’s protocol. Here, PicoGreen^™^ was diluted in 10 mM Tris-HCL, 1 mM EDTA with pH 7.5 (TE). A DNA stock solution (100 μg/mL) was dissolved in TE solution and serial dilution from 0 to 200 ng of DNA/mL was performed to establish a calibration curve. Afterwards, 10 μL of each standard and sample were added in triplicate to a black flat bottomed 96 well plate and further 190 ul of diluted PicoGreen^™^ in TE solution were added. Fluorescence emission was read at 520nm using an excitation wavelength of 480nm in a plate reader with 0.93 seconds per well of dwell time.

### Statistics

Statistical analysis was performed using GraphPad Prism 9 (GraphPad Software, USA). Where appropriate, one-way analysis of variance (ANOVA) or two-way ANOVA were used as statistical tests followed by Tukey’s multiple comparison. Results are presented as mean ± standard deviation and differences are considered as statistically significant for p-value ≤ 0.05.

## Acknowledgments

NN is supported by a Trinity College Dublin, Provost’s PhD Award, and the TCD FLIM core unit directed by MM is supported by a SFI Infrastructure Programme: Category D Opportunistic Funds Call (16/RI/3403). This work was also partially supported by EPSRC and SFI Centre for Doctoral Training in Engineered Tissues for Discovery, Industry and Medicine, Grant Number EP/S02347X/1 (to MM and MS) and in part by a grant from Science Foundation Ireland (SFI) and the European Regional Development Fund (ERDF) under grant number 13/RC/2073_P2. The authors acknowledge funding from Science Foundation Ireland (SFI) Frontiers for the Future Project Grant (19/FFP/6533).

## Author contributions

Conceptualization, Data curation, Software, Formal analysis, Validation, Investigation, Visualization, Methodology, Writing – original draft, Writing – review and editing were performed by MM and NN. MS contributed towards qPCR execution and analysis and writing of the paper. DH contributed towards hypothesis development and editing the final draft of the manuscript.

