## Supplemental for "2P-FLIM unveils time-dependent metabolic shifts during osteogenic differentiation with a key role of lactate to fuel osteogenesis via glutaminolysis identified"

Supplemental Table 1 – Gene symbol, name, accession number, unique assay ID and amplicon length of genes used for qPCR

| Gene Symbol | Gene Name | RefSeq Accession No | Unique Assay Id | Amplicon Length |
|---|---|---|---|---|
| ALPL | alkaline phosphatase, | NC_000001.10,NG_008940.1,NT_004610.19 | qHsaCID0010031 | 99 |
| GOT1 | glutamic-oxaloacetic transaminase 1 | NC_000010.10,NT_030059.13 | qHsaCID0006686 | 140 |
| SLC16A3 | monocarboxylic acid transporter 4 | NC_000017.10,NT_010663.15 | qHsaCID0014322 | 180 |
| BGLAP | bone gamma-carboxyglutamate (gla) protein | NC_000001.10,NT_004487.19 | qHsaCED0038437 | 69 |
| GOT2 | glutamic-oxaloacetic transaminase 2, mitochondrial | NC_000016.9,NT_010498.15 | qHsaCID0013583 | 152 |
| P4HA1 | prolyl 4-hydroxylase | NC_000010.10,NT_030059.13 | qHsaCID0009945 | 148 |
| SLC16A4 | monocarboxylic acid transporter 5 | NC_000001.10,NT_032977.9 | qHsaCED0036763 | 78 |
| P4HA2 | prolyl 4-hydroxylase | NC_000005.9,NT_034772.6 | qHsaCID0012216 | 79 |
| SOX9 | SRY (sex determining region Y)-box 9 | NC_000017.10,NG_012490.1,NT_010783.15 | qHsaCED0021217 | 77 |
| LDHA | lactate dehydrogenase A | NC_000011.9,NG_011816.1,NT_009237.18,NG_008185.1 | qHsaCED0056429 | 68 |
| P4HA3 | prolyl 4-hydroxylase, | NC_000011.9,NT_167190.1 | qHsaCID0009995 | 104 |
| SPP1 | secreted phosphoprotein 1 | NC_000004.11,NT_016354.19 | qHsaCID0012060 | 99 |
| GLS | glutaminase | NC_000002.11,NT_005403.17 | qHsaCID0007574 | 120 |
| PTGS2 | prostaglandin-endoperoxide synthase 2 | NC_000001.10,NT_004487.19 | qHsaCID0020933 | 148 |
| UEVLD | UEV and lactate/malate dehyrogenase domains | NC_000011.9,NG_012138.1,NT_009237.18 | qHsaCED0036634 | 75 |
| GLS2 | glutaminase 2 mitochondrial | NC_000012.11,NT_029419.12 | qHsaCED0002029 | 72 |
| RUNX2 | runt-related transcription factor 2 | NC_000006.11,NT_007592.15,NG_008020.1 | qHsaCID0006726 | 80 |
| GLUD1 | glutamate dehydrogenase 1 | NC_000010.10,NG_013010.1,NT_030059.13 | qHsaCED0038578 | 73 |
| LDHB | lactate dehydrogenase B | NC_000012.11,NT_009714.17,NG_017038.1 | qHsaCID0012068 | 60 |
| SLC16A1 | monocarboxylic acid transporter 1 | NC_000001.10,NT_032977.9,NG_015880.1 | qHsaCID0008777 | 64 |
| GLUD2 | glutamate dehydrogenase 2 | NC_000023.10,NG_016456.1,NT_011786.16 | qHsaCED0038220 | 102 |
| LDHC | lactate dehydrogenase C | NC_000011.9,NG_011816.1,NT_009237.18 | qHsaCID0018614 | 130 |
| SLC16A2 | monocarboxylic acid transporter 8 | NC_000023.10,NG_011641.1,NT_011669.17 | qHsaCID0015666 | 149 |

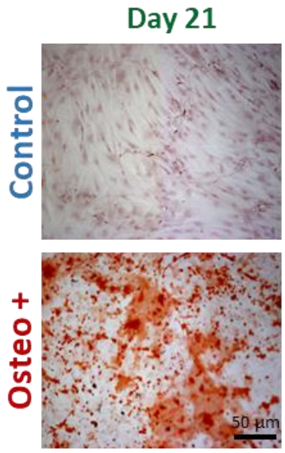


Supplemental Figure 1 - Alizarin red staining of hMSCs after 21 days of incubation in either Xpan or Osteo+ cell culture media.

Supplemental Table 2 – Mahalanobis distance, Hotelling $T^2$ stats, F-value, Critical F-Values, P-value and significance results of PCA statistical significance analysis

| | Mahalanobis Distance | Two-Sample T2 Stats | F-Value | Critical F-Value | P-value | Significance |
|---|---|---|---|---|---|---|
| Osteo vs Xpan | 4.121 | 25.478 | 9.554 | 9.552 | 0.025 | YES |
| Figure 4.3 | | | | | | |
| Osteo vs Xpan (D0) | 1.165 | 8.151 | 3.89 | 3.467 | 0.2063 | NO |
| Osteo vs Xpan (D3) | 1.986 | 23.665 | 11.295 | 3.467 | 0.105 | NO |
| Osteo vs Xpan (D7) | 3.223 | 62.314 | 29.748 | 3.467 | 0.044 | YES |
| Osteo vs Xpan (D14) | 5.32 | 169.823 | 81.052 | 3.467 | 0.012 | YES |
| Figure 4.5 | | | | | | |
| Osteo vs Xpan | 3.105 | 49.152 | 11.264 | 9.552 | 0.048 | YES |
| Xpan vs Lact | 11.06 | 183.472 | 68.802 | 9.552 | 0.0005 | YES |
| Lact vs Osteo | 6.4905 | 63.19 | 23.696 | 9.552 | 0.006 | YES |
| Figure 4.6 | | | | | | |
| Xpan vs Osteo | 5.569 | 139.584 | 65.431 | 3.682 | 0.01 | YES |
| Osteo vs Lact | 3.127 | 43.992 | 20.621 | 3.682 | 0.0472 | YES |
| Lact vs Xpan | 3.888 | 68.019 | 31.884 | 3.682 | 0.0289 | YES |
| Figure 4.8 | | | | | | |
| Osteo vs Xpan | 7.513 | 84.666 | 31.75 | 9.552 | 0.0034 | YES |
| Xpan vs Lact | 5.232 | 41.059 | 15.397 | 9.552 | 0.0127 | YES |
| Lact vs Osteo | 4.942 | 36.641 | 13.74 | 9.552 | 0.0151 | YES |

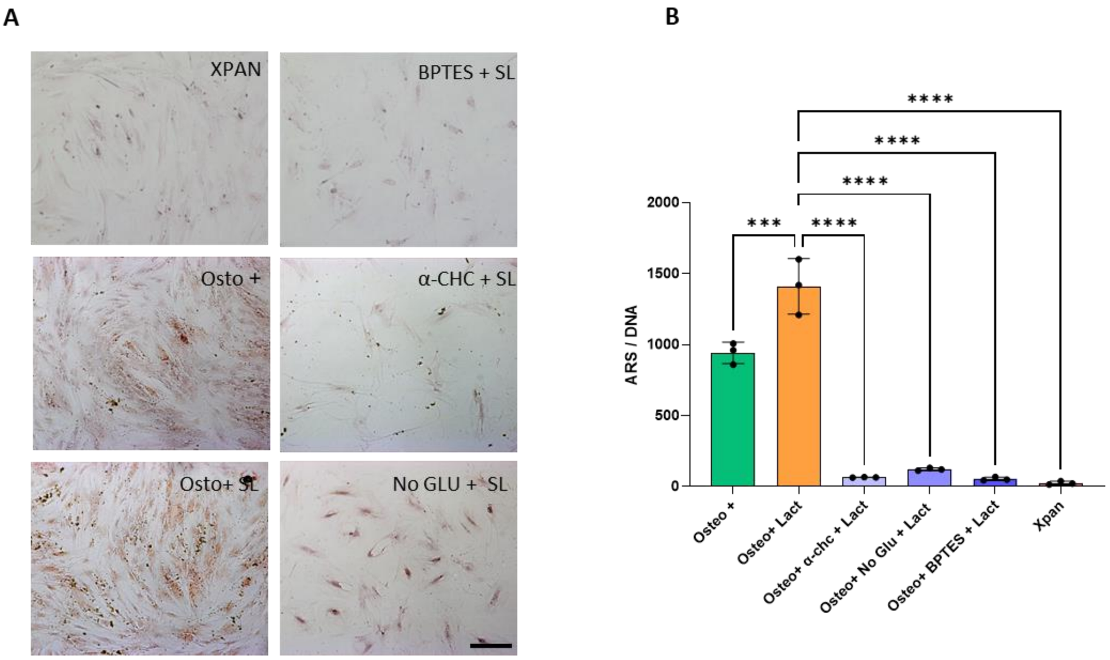


Supplemental Figure 2 – Supplementation of Osteo + cell culture medium with lactate and metabolic inhibitors. (A) Alizarin red staining of hMSCs after 14 days of cell culture in several cell culture medium formulations. (B) Alizarin red quantification per DNA of hMSCs after 14 days of cell culture
